# IgG-aggregates rapidly up-regulate FcgRI expression at the surface of human neutrophils in a FcgRII-dependent fashion: A crucial role for FcgRI in the generation of reactive oxygen species

**DOI:** 10.1101/2020.03.04.975094

**Authors:** Sandrine Huot, Cynthia Laflamme, Paul R. Fortin, Eric Boilard, Marc Pouliot

**Affiliations:** Département de microbiologie et immunologie, Faculté de Médecine de l’Université Laval, Centre de Recherche du CHU de Québec-Université Laval, Québec City, QC, Canada; Division de Rhumatologie, Département de Médecine, CHU de Québec-Université Laval, Québec City, QC, Canada; Axe maladies infectieuses et immunitaires, Centre de Recherche du CHU de Québec-Université Laval, Québec City, QC, Canada

## Abstract

Autoimmune complexes are an important feature of several autoimmune diseases such as lupus, as they contribute to tissue damage through the activation of immune cells. Neutrophils, key players in lupus, interact with immune complexes through Fc gamma receptors (FcgR). Incubation of neutrophils with aggregated-IgGs caused degranulation and increased the surface expression of FcgRI within minutes in a concentration-dependent fashion. After 30 min, IgG aggregates (1 mg/ml) up-regulated FcgRI by 4.95 ± 0.45-fold. FcgRI-positive neutrophils reached 67.24% ± 6.88% on HA-IgGs stimulated neutrophils, from 3.12% ± 1.62% in non-stimulated cells, ranking IgG-aggregates among the most potent known agonists. FcgRIIa, and possibly FcgRIIIa, appeared to mediate this up-regulation. Also, FcgRI-dependent signaling proved necessary for reactive oxygen species (ROS) production in response to IgG-aggregates. Finally, combinations of bacterial materials with aggregates dramatically boosted ROS production. This work suggests FcgRI as an essential component in the response of human neutrophils to immune complexes leading to the production of ROS, which may help explain how neutrophils contribute to tissue damage associated with immune complex-associated diseases, such as lupus.

## Introduction

Immune complex diseases, such as rheumatoid arthritis and systemic lupus erythematosus, encompass a diverse group of inflammatory conditions characterized by antigen-antibody deposition. Typical manifestations are glomerulonephritis, synovitis, and dermal vasculitis (1). Activation of neutrophils, resulting from the interaction with immune complexes, leads to the production of oxygen-free radicals and the release of lysosomal enzymes (2), which may bring about tissue inflammation and organ damage.

Phagocytes, including neutrophils, bind to immune complexes with their Fc gamma receptors (FcgRs), serving as molecular links between antibody and immunological responses (3). Neutrophils express the transmembrane FcgRIIa (CD32a) and FcgRIIIb (CD16b), expressed at 10-fold higher density than FcgRIIa [Reviewed in (4)]. FcgRIIb is also present, albeit to a lesser degree (4). Both FcgRIIIb and FcgRIIa are low-affinity receptors, exclusive to higher primates, and displaying specialized context-dependent functions in neutrophil recruitment (5, 6). Unique to neutrophils, FcgRIIIb is the only Fc receptor anchored by a glycosylphosphatidylinositol (GPI)-linker to the plasma membrane, and lacking a cytoplasmic domain (7). Despite the absence of a known molecular adaptor, multiple studies have reported that FcgRIIIb can induce cellular responses such as Ca^2+^ mobilization (8), nuclear factor activation (9), removal of immune complexes (10) and NET formation (11). Recently, the expression of FcgRIIIa (CD16a) was demonstrated in neutrophils (12). FcgRIIIa and FcgRIIIb share 96% sequence identity in their extracellular IgG-binding regions (7). FcgRIIIa is a classic activating receptor that, upon stimulation, associates with an Immunoreceptor Tyrosine-based Activation Motif (ITAM)-signaling system. Thus, FcgRIII signaling in neutrophils is still not completely understood. On the other hand, FcgRIIa, with its integrated signaling ITAM, has been identified as an essential mediator of destructive antibody-based inflammation in autoimmunity (5) and is necessary for neutrophil recruitment within glomerular capillaries following IgG deposition (13, 14).

FcgRI (CD64) is the only high-affinity receptor of immunoglobulin G (IgG) in humans (15). Like FcgRIIIa, FcgRI is an ‘activator’ using an intracellular ITAM-mediated signaling system to promote phagocytosis, respiratory burst, antibody-dependent cell-mediated cytotoxicity, cytokine production, and NETosis (3, 16, 17). Nonetheless, FcgRI surface expression is negligible in circulating healthy individuals’ neutrophils and is stored in intracellular vesicles (18). Modulation of FcgRI by cytokines or growth factors was reported more than three decades ago. IFN-gamma increased FcgRI on neutrophils after 18 hrs of incubation (19). *In vivo*, but not *in vitro* application of granulocyte-colony stimulating factor (G-CSF) increased FcgRI expression on neutrophils within days, and potentiated their tumor cell killing capabilities (20, 21). Also, neutrophil FcgRI expression increases in complications associated with bacterial infections (22-24). Currently, neutrophil FcgRI serves as a sensitive diagnostic marker for early-onset neonatal infections (25).

While monocyte FcgRI expression is positively associated with immune inflammation and lupus (26-28), its expression on circulating neutrophils is barely elevated in patients with lupus, unless an infection is present (29). Moreover, the implication of neutrophil FcgRI in responses to immune complexes is not documented.

In this paper, we investigated the impact of aggregated-IgGs, used as a model of immune complexes, on neutrophil activation. We show that the engagement of FcgRs by IgG-aggregates up-regulates FcgRI at the surface of human neutrophils from intracellular stores. Moreover, we assessed the necessity of FcgRI activation for downstream cellular responses. This work reveals FcgRI as an essential component for ROS production following the engagement of FcgRs.

## Materials & Methods

### Materials

Dextran-500, Lipopolysaccharide (LPS from *Escherichia coli* 0111: B40), Actinomycin D (from *Streptomyces* sp.), Formyl-methionyl-leucyl phenylalanine (fMLP), Wortmannin (from *Penicillium fumiculosum*), Phorbol 12-myristate 13-acetate (PMA), Luminol, Isoluminol, Cycloheximide and SB216763 were purchased from Sigma-Aldrich (Oakville, ON, Canada). Lymphocyte separation medium was purchased from Wisent (St-Bruno, QC, Canada). AS605240, U0126, and U73122 were obtained from Cayman Chemical (Ann Arbor, MI, USA). PRT-060318 was obtained from Selleckchem (Houston, TX, USA). Gö6976 and PP-2 were purchased from Calbiochem (San Diego, CA, USA). Recombinant human granulocyte (G), granulocyte-macrophage (GM) colony-stimulating factor (CSF), interferon (IFN)-gamma (γ), and tumor necrosis factor (TNF) were purchased from PeproTech (Rocky Hill, NJ, USA). Nexinhib20 was obtained from Tocris (Oakville, ON, Canada). IFN-alpha (α) was purchased from PBL Assay Science (Piscataway, NJ, USA). Human immunoglobulins G (IgGs): Lyophilized human IgGs, purified from human plasma or serum by fractionation (purity of greater than 97% as determined by SDS-PAGE analysis) were purchased from Innovative Research, Inc. (Novi, MI, USA). Phosphatidylinositol-Specific Phospholipase C (PI-PLC), 7-Aminoactinomycin D (7-AAD) and SYTOX green were purchased from Thermo Fisher Scientific (Burlington, ON, Canada). Cytochrome C was obtained from Bio Basic (Markham, ON, Canada). Human MPO and human Lactoferrin ELISA kits were purchased from Assaypro (St. Charles, MO, USA). MMP-9 Duoset ELISA was obtained from R&D Systems, Inc. (Oakville, ON, Canada). A list of compounds, molecular targets, concentrations, and solvents, is presented in **Table 2**. A list of agonists and concentrations is presented in **Table 3**.

**Table 1.** Impact of heat-aggregated (HA)-IgGs on neutrophil surface markers. Human neutrophils were incubated with HA-IgGs (1 mg/ml, 30 min, 37°C), and surface markers were monitored by flow cytometry, as described in *Methods*. Markers of the following parameters were measured: maturity and adhesion (CD10, CD15); adhesion and migration (total, and activated form (^act^) of CD11b); degranulation of primary (CD63), secondary (CD66b, CD11b), tertiary, and secretory vesicles (CD11b); Fc gamma receptors FcgRI (CD64), FcgRII (CD32), and FcgRIII (CD16). FcgRI and CD63 were most up-regulated (in Bold), both in a comparable fashion. For each marker, results are expressed as the ratio obtained from mean fluorescence intensity after incubations with HA-IgGs, relative to non-stimulated cells. Mean ± SEM (n=4 experiments). *p<0.05

| Marker | Relative Fluorescence Intensity | p value |
| --- | --- | --- |
| CD10 | 1.69±0.16 * | 0.037 |
| CD15 | 1.48±0.13 * | 0.048 |
| CD11b | 1.29±0.08 * | 0.023 |
| CD11b<sup>act</sup> | 1.61±0.21 * | 0.045 |
| CD66b | 1.02±0.09 | 0.884 |
| **CD63** | **3.99±0.55 \*** | **0.001** |
| **FcgRI (CD64)** | **4.61±0.79 \*** | **0.009** |
| FcgRII (CD32) | 0.65±0.11* | 0.004 |
| FcgRIII (CD16) | 0.79±0.14 * | 0.048 |

**Table 2.** List of compounds, their molecular target, concentrations, and diluting buffers.

| Compound | Molecular target | Final conc. | Solvent |
| --- | --- | --- | --- |
| Cycloheximide | Protein synthesis | 10 µg/ml | EtOH |
| Actinomycin D | Transcription | 2 µg/ml | DMSO |
| PP-2 | Src family of kinase | 10 µM | DMSO |
| Wortmannin | PI3K | 200 nM | DMSO |
| AS605240 | PI3K | 10 µM | DMSO |
| U73122 | PLC | 10 µM | DMSO |
| Gö6976 | PKCa, PKCb<sub>1</sub> | 1 µM | DMSO |
| U0126 | MEK1/2 | 10 µM | DMSO |
| PRT-060318 | Syk | 10 µM | HBSS |
| SB216763 | GSK-3α | 10 µM | DMSO |
| Nexinhib20 | Rab27a-JFC1 interaction | 10 µM | DMSO |
| 10.1 | FcγRI (CD64) | 3 µg/ml | Aqueous buffer |
| IV.3 | FcγRIIa (CD32a) | 2.5 µg/ml | Aqueous buffer |
| 3G8 | FcγRIII (CD16) | 4 µg/ml | Aqueous buffer |
| 11.5 | FcγRIIIb (CD16b) | 2.5 µg/ml | Aqueous buffer |
| PI-PLC | FcγRIIIb (CD16b) | 0.5 U/ml | Aqueous buffer |
| Fc-Block | Fc receptors | 2.5 mg/100 µl | Aqueous buffer |

**Table 3.** List of agonists and final concentrations.

| Agonist | Final conc. |
| --- | --- |
| TNF | 100 ng/ml |
| GM-CSF | 1.4 nM |
| G-CSF | 1.4 nM |
| IFNα | 1000 U/ml |
| IFNγ | 100 ng/ml |
| LPS | 1 μg/ml |
| fMLP | 100 nM |
| PMA | 10 nM |

### Antibodies

Purified mouse anti-human FcgRIII (3G8), Alexa Fluor®-labeled 647 mouse anti-human CD16 (3G8), PE-labeled mouse anti-human FcgRIIIb (Clone CLB-gran11.5), purified mouse anti-human FcgRI (10.1), V450-labeled mouse anti-human FcgRI (10.1) and PE-labeled mouse anti-human FcgRII (Clone FLI8.26, also known as 8.26), were purchased from BD Biosciences (San Jose, CA, USA). Monoclonal anti-human FcgRIIa (IV.3) was obtained from Bio X Cell (West Lebanon, NH, USA). This blocking antibody recognizes a native extracellular epitope of FcgRIIa (30).

### Ethics

All experiments involving human tissues received approval from the Université Laval ethics committee.

### Isolation of human neutrophils

Informed consent was obtained in writing from all donors. Data collection and analyses were performed anonymously. Neutrophils were isolated as originally described (31) with modifications (32). Briefly, venous blood collected on isocitrate anticoagulant solution from healthy volunteers was centrifuged (250 x *g*, 10 min), and the resulting platelet-rich plasma was discarded. Leukocytes were obtained following sedimentation of erythrocytes in 2% Dextran-500. Neutrophils were then separated from other leukocytes by centrifugation on a 10 ml lymphocyte separation medium. Contaminating erythrocytes were removed using 20 sec of hypotonic lysis. Purified granulocytes (> 95% neutrophils, < 5% eosinophils) contained less than 0.2% monocytes, as determined by esterase staining. Viability was greater than 98%, as determined by trypan blue dye exclusion. The entire cell isolation procedure was carried out under sterile conditions at room temperature.

### Cell incubations

Neutrophils were resuspended at a concentration of 10 x 10^6^ cells/ml at 37°C in Hank’s Balanced Salt Solution (HBSS) containing 10 mM HEPES pH 7.4, 1.6 mM Ca^2+^ and no Mg^2+^, 10% human serum, and supplemented with 0.1 U/ml adenosine deaminase (ADA) to prevent the accumulation of endogenous adenosine in the medium, thus minimizing the previously demonstrated modulating effects of adenosine on inflammatory factor production by neutrophils (33-35).

### Preparation of heat-aggregated IgGs

Heat-aggregated (HA)-IgGs were essentially prepared as originally described (36), with modifications. Briefly, soluble aggregates were prepared daily by resuspending IgGs in HBSS 1X at a concentration of 25 mg/ml, and heating at 63°C for 75 min.

### Flow Cytometry

Following appropriate treatments, cell suspensions were spin and resuspended in HBSS containing human Fc-Block (BD Biosciences). Cells were incubated with V450-labeled mouse anti-human FcgRI (CD64), Alexa Fluor®-labeled 647 mouse anti-human FcgRIII (CD16) and PE-labeled mouse anti-human FcgRII (CD32) for 30 min in the dark. HBSS and 7-AAD (viability) were added (400 μl), and samples were analyzed using a FACS Canto II flow cytometer with FACSDiva software, version 6.1.3 (BD Biosciences). Cell gating strategy and representative data are presented in **Supplemented Figure S1**.

### Viability

Neutrophil viability was assessed using a FITC Annexin V Apoptosis Detection Kit (BD Biosciences). Briefly, cell pellets were suspended in 100 μL of binding buffer. Annexin V and propidium iodide (5 µl each) were added to each sample. After 15 min, 400 μL of binding buffer was added, and samples were analyzed by flow cytometry. Gating was determined using control samples labeled individually with either Annexin V or propidium iodide.

### Degranulation. A) Surface markers

Neutrophils were processed for flow cytometry as described above. Surface levels of: CD63 (primary granules); CD66b (secondary granules), CD11b, (secondary, tertiary granules, and secretory vesicles); CD10, and CD35 (secretory vesicles) were monitored (37). *B) Matrix proteins*. Cell-free supernatants were stored at –20°C until their analysis for myeloperoxidase (MPO), lactoferrin, and gelatinase (MMP-9) content, with commercially-available ELISAs, according to the manufacturers’ instructions.

### NET production

Performed essentially as in (38). Briefly, neutrophils (5 x 10^4^ cells/well) in HBSS containing HEPES (10 mM), Ca^2+^ (1.6 mM) and supplemented with ADA (0.1 U/ml) were aliquoted (200 µl) into 96 well plates and left to settle for 30 min at 37°C. When present, Nexinhib20 or 10.1 antibody was added 30 min before adding HA-IgGs, or PMA (10 nM) as a positive control for NET induction (38). Plates were incubated at 37°C (5% CO_2_) for up to 4 hrs; SYTOX green (5 μM final concentration), a cell-impermeable nucleic acid stain, was added at indicated times and NET formation was evaluated by measuring the fluorescence (excitation/emission: 504/523 nm) in each well after subtraction of the background fluorescence.

### Cytokine/chemokine release

Cell-free supernatants were stored at –20°C until their analysis for chemokine/cytokine content with a multiplexed bead-based immunoassay (BD(tm) Cytometric Bead Array), using FCAP Array software version 3.0 (BD Biosciences). CXCL8 (IL-8), CCL2 (MCP-1), CCL3 (MIP-1alpha), TNF, IL-1alpha, IL-1beta, IL-6, and IFN-alpha were monitored.

### Production of reactive oxygen species

Reactive oxygen species (ROS) production was measured as previously described (39) with modifications. Briefly, neutrophils were resuspended at 1 x 10^6^ cells/ml in the presence of either 2.5% membrane non-permeable cytochrome c (v/v), 10 µM of membrane-permeable luminol, or 10 µM of membrane non-permeable isoluminol. Neutrophils (200 µl) were placed in 96-well microplates, treated as indicated, and stimulated with 1 mg/ml of free IgGs or 1 mg/ml of HA-IgGs. Suspensions were incubated at 37°C in the microplate reader Infinite M1000 PRO with i-control 2.0 software (Tecan, Morrisville, NC, USA). Luminescence intensity (luminol and isoluminol) and optical density (cytochrome C; 550 nm with a correction at 540 nm) were monitored every 5 min. The amount of superoxide anion produced in the cytochrome C reduction assay was calculated using the formula published by Dahlgren and Karlsson (40).

### Statistical analysis

Statistical analysis was performed using GraphPad PRISM version 8 (GraphPad Software, San Diego, CA, USA). Where applicable, values are expressed as the mean ± standard error of the mean (SEM). Statistical analysis was performed using a two-tailed Student’s t-test (paired comparisons). Differences were considered significant (marked by an asterisk*) when p < 0.05.

## Results

### Heat-aggregated IgGs rapidly up-regulate FcgRI expression on neutrophils

In order to gain insights about the implication of neutrophils in inflammatory events resulting from interactions with immune complexes, we incubated human neutrophils with heat-aggregated (HA)-IgGs and measured the surface expression of activation markers by flow cytometry. Such aggregates are a good initial model of immune complexes, as they cause no bias based on the antigen (41). HA-IgGs significantly increased markers associated with maturity (CD10) (42) and adhesion (CD11b, CD15) (43, 44) (**Table 1**). CD11b is present on secondary, tertiary granules and secretory vesicles (45), thereby suggesting their exocytosis as well, although CD66b, expressed in secondary granules, was not affected. HA-IgGs had the strongest and comparable impact on FcgRI (CD64) and primary granule marker CD63 (Relative fluorescence intensities of 4.61±0.79 and 3.99±0.55, respectively). In contrast, HA-IgGs decreased the expression of FcgRII (CD32), as reported earlier (46), and FcgRIII (CD16). Such an up-regulation of neutrophil-FcgRI is unprecedented and warrants further investigation.

HA-IgGs increased FcgRI in a concentration-dependent and saturable fashion (**Fig. 1A**). At the highest concentration, the proportion of FcgRI-positive neutrophils reached 75%, from 2.5% on resting cells (**Fig. 1C**). HA-IgGs up-regulated the surface expression of FcgRI within minutes and remained elevated for at least 60 min (**Fig. 1B, D**). Free IgGs had little to no effect on FcgRI expression. Conversely, HA-IgGs decreased the signal intensity of FcgRII and FcgRIII, by 63% and 47%, respectively, compared to non-stimulated cells (**Fig. 1E, I**), but did not affect percentages of positive cells (**Fig. 1G, H, K, L**). Overall, the impact of HA-IgGs on FcgRs expression was potent and occurred within minutes (**Fig. 1B, F, J**).

**Figure 1.**
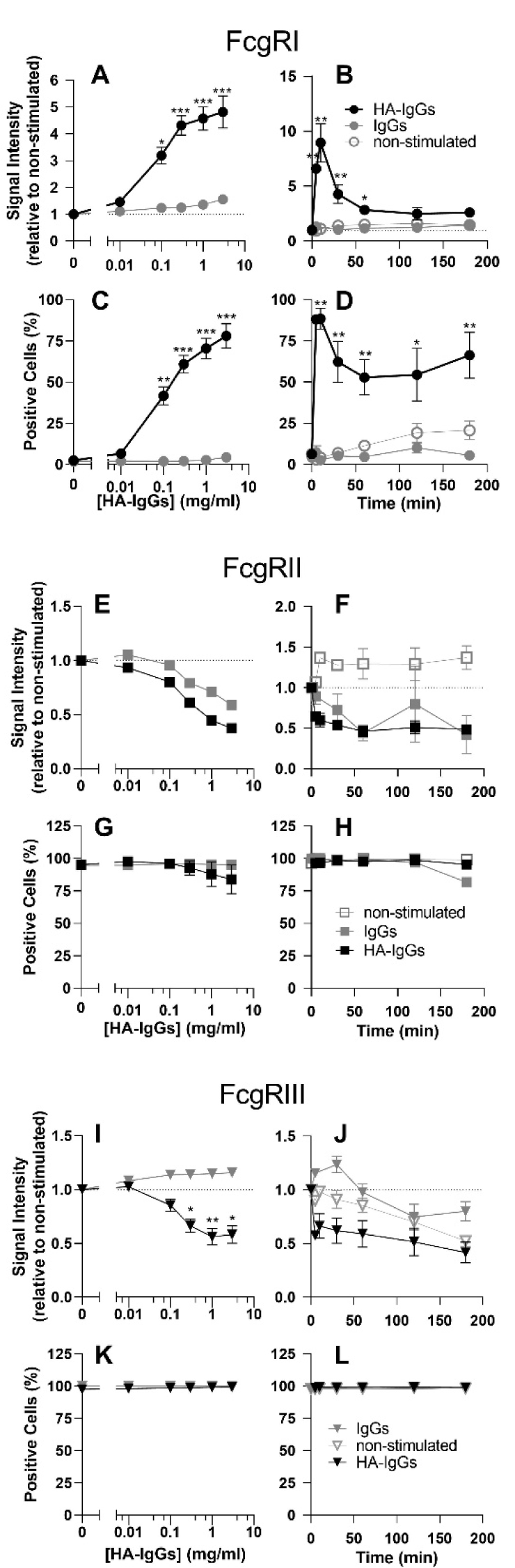
HA-IgGs rapidly up-regulate FcgRI on human neutrophils, in a concentration-dependent fashion. Neutrophils were incubated with indicated concentrations of heat-aggregated (HA)-IgGs, or free IgGs, for 30 min at 37°C (left panels), or for indicated times at 1 mg/ml (right panels). Surface expressions of FcgRI (**A-D**), FcgRII, (**E-H**), and FcgRIII (**I-L**) were measured by flow cytometry. Results are expressed as the fluorescence signal intensity relative to non-stimulated cells and are from n=4 experiments, each performed with neutrophils from different healthy donors. *p<0.05, **p<0.01, ***p<0.001

### HA-IgGs are a strong stimulus for the up-regulation of FcgRI on human neutrophils

Injections of IFN-gamma or G-CSF have been reported to up-regulate FcgRI on neutrophils, upon lengthy exposures (19-21). We incubated neutrophils with LPS, IFN-alpha, IFN-gamma, fMLP, TNF, G-CSF, or GM-CSF, alone or in combination with HA-IgGs, and measured FcgRs expression. By themselves, none of these factors had any significant effect on FcgRI expression within 30 min (**Fig. 2A, white bars**), while HA-IgGs led to a 4.9-fold increase (n=4) in FcgRI mean fluorescence intensity (**Fig. 2A, black bars**); FcgRI-positive neutrophils reached 67% on HA-IgGs stimulated neutrophils, from 3% in non-stimulated cells (**Fig. 2B**). When combined with HA-IgGs, LPS and GM-CSF had significantly potentiating effects on FcgRI surface expression. While all factors tended to increase percentages of FcgRI-positive neutrophils, significance was not reached. GM-CSF slightly increased the basal expression of FcgRIII (**Fig. 2D**); LPS and GM-CSF potentiated the decreasing-effect of HA-IgGs on FcgRIII. Finally, none of these factors affected FcgRII expression (**Fig. 2C**). Overall, in these conditions, HA-IgGs constitute a potent stimulus for FcgRI up-regulation.

**Figure 2.**
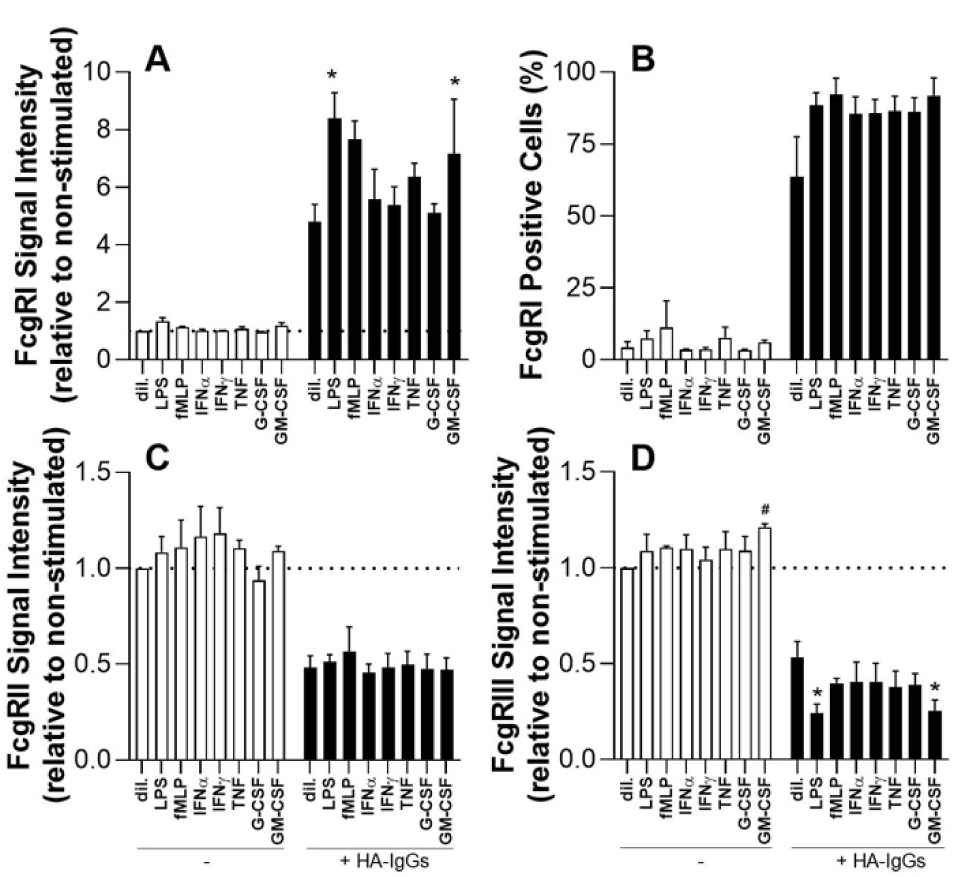
HA-IgGs are a potent stimulus for FcgRI up-regulation in human neutrophils. Cells were incubated with indicated agonists or diluent (dil.) for 15 min (37°C), then with HA-IgGs for 30 min. FcgRs expression was determined by flow cytometry. **A, C, D)** For each of the FcgR, results are expressed as the fluorescence signal intensity relative to non-stimulated cells (dil., white bars). **B)** Percentages of FcgRI-positive neutrophils. From n=4 experiments, each performed with neutrophils from different donors. Mean ± SEM. *Statistically different from HA-IgGs alone (dil., black bar), or ^#^from resting cells. p<0.05

### FcgRIIa, but not FcgRIIIb, is instrumental in the up-regulation of FgcRI by HA-IgGs

We initially assessed the possible implication of FcgRIII in the up-regulation of neutrophil-FcgRI by HA-IgGs. The FcgRIII-blocking monoclonal antibody 3G8 did not prevent FcgRI up-regulation. Moreover, the FcgRIIIb-specific monoclonal antibody 11.5 significantly enhanced FcgRI up-regulation, so did the shedding of FcgRIIIb by PI-PLC (47) (**Fig. 3A**), arguing against a significant role for FcgRIIIb in FcgRI up-regulation, but leaving the possibility open for FcgRIIIa.

**Figure 3.**
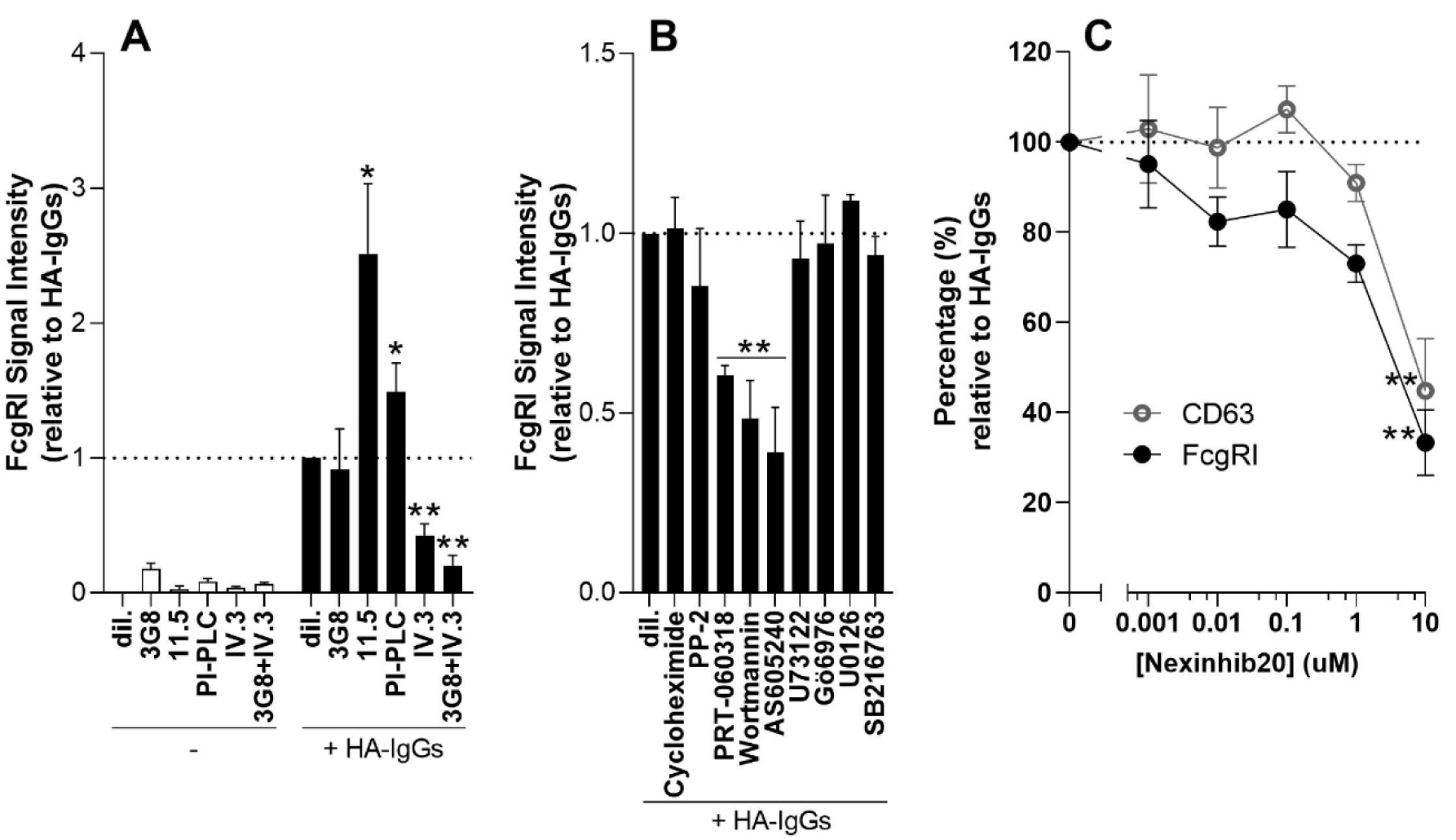
Signaling pathways involved in the up-regulation by HA-IgGs. **A)** Neutrophils were incubated with the following whole antibodies: 3G8 (FcgRIII), 11.5 (FcgRIIIb), IV.3 (FcgRII), or with PI-PLC for 15 min (37°C), then stimulated with HA-IgGs for an additional 30 min. **B)** Neutrophils were incubated with indicated pharmacological inhibitors before stimulation with HA-IgGs. FcgRI surface expression was measured by flow cytometry. Results are expressed as the fluorescence signal intensity relative to HA-IgGs alone (dil., black bar),. **C)** Neutrophils were incubated with indicated concentrations of Nexinhib20 for 15 min before stimulation with HA-IgGs. Mean ± SEM, n=3. Significantly higher*, and significantly lower** than HA-IgGs alone, p<0.05.

As expected, the FcgRIIa-blocking monoclonal antibody IV.3 effectively prevented the up-regulation of FcgRI, by approximately 60%. Simultaneous blockade of FcgRIII and FcgRIIa did not result in more inhibition. Together, these results confirm a role for FcgRIIa, but not for FcgRIIIb, in FcgRI up-regulation.

### Syk and PI3K mediate the up-regulation of FcgRI by HA-IgGs

Both FcgRIIa and FcgRIIIa are activators, signaling either through an intrinsic ITAM (FcgRIIa), or through association with an ITAM-containing gamma subunit (FcgRIIIa) (48). Tyrosine residues in the ITAM, phosphorylated by Src-family protein kinases, constitute anchoring sites for the spleen tyrosine kinase (Syk), the activation of which leads to stimulation of phosphatidylinositol 3-kinase (PI3K), phospholipase Cγ (PLCγ), ERK (extracellular signal-regulated kinase), and other downstream pathways (49). Despite an inhibitory trend, the Src-family kinase inhibitor PP-2 only had a modest impact on FcgRI up-regulation (**Fig. 3B**). In contrast, the Syk inhibitor PRT-060318, and two structurally distinct PI3K inhibitors, Wortmannin and AS605240, each prevented the up-regulation of FcgRI, approximately by half. Other downstream pathways, including PLC (U73122), PKC (Gö6976), MEK/ERK (U0126), or glycogen-synthase-kinase (GSK)-3 (SB216763), did not appear to be involved. Finally, the protein synthesis inhibitor cycloheximide also had no sizeable impact.

### FcgRI is pre-formed, stored in intracellular granules

The rapid up-regulation of FcgRI suggested granule localization for FcgRI. To confirm, we used the neutrophil exocytosis inhibitor Nexinhib20. As shown in **Figure 3C**, Nexinhib20 effectively reversed FcgRI increase, in a concentration-dependent fashion and up to 66±7%, at 10 µM. The effect of Nexinhib20 on CD63 was comparable. With a rapid, protein-synthesis independent up-regulation by HA-IgGs and a profile of expression following that of CD63, these results support a primary granule localization for pre-formed FcgRI, in resting neutrophils.

### HA-IgGs induce degranulation, MCP-1 release, and intracellular ROS production

Given this robust and rapid up-regulation in FcgRI surface expression, we evaluated the functional implication of FcgRI in some meaningful neutrophil responses, namely: cell viability, degranulation, NET formation, cytokine release, and ROS production. HA-IgGs did not decrease neutrophil viability (**Suppl. Fig. S2**); in fact, HA-IgGs delayed early apoptosis after 24 hrs and 48 hrs. The blockade of FcgRI with the FcgRI-specific 10.1 antibody had no significant impact. In degranulation experiments, HA-IgGs provoked significant, rapid release of MPO, lactoferrin, and MMP-9 in cell-free supernatants (**Suppl. Fig. S3**); FcgRI blockade had no inhibitory effect. In contrast, Nexinhib20 completely inhibited degranulation. HA-IgGs did not induce NET formation (**Suppl. Fig. S4**). Regarding the release of cytokines, HA-IgGs specifically stimulated the release of CCL-2 (MCP-1) (**Suppl. Fig. S5**). This observation is of interest because in the serum and urine of patients with lupus, MCP-1 levels are elevated compared to healthy controls; moreover, those levels positively correlate with lupus activity (50). Here also, FcgRI blockade did not reverse the HA-IgG-triggered release of MCP-1.

Also, we evaluated the production of ROS elicited by HA-IgG-stimulated neutrophils, by monitoring: the overall MPO-dependent production of HOCl with luminol, the extracellular-only MPO-dependent production of HOCl with isoluminol, and the extracellular-only release of O_2_ ^-^ reactive species with cytochrome C. As shown in **Figure 4**, the incubation of neutrophils with HA-IgGs resulted in the intracellular-only generation of ROS, as reported earlier (39). Free-IgGs had little to no effect.

**Figure 4.**
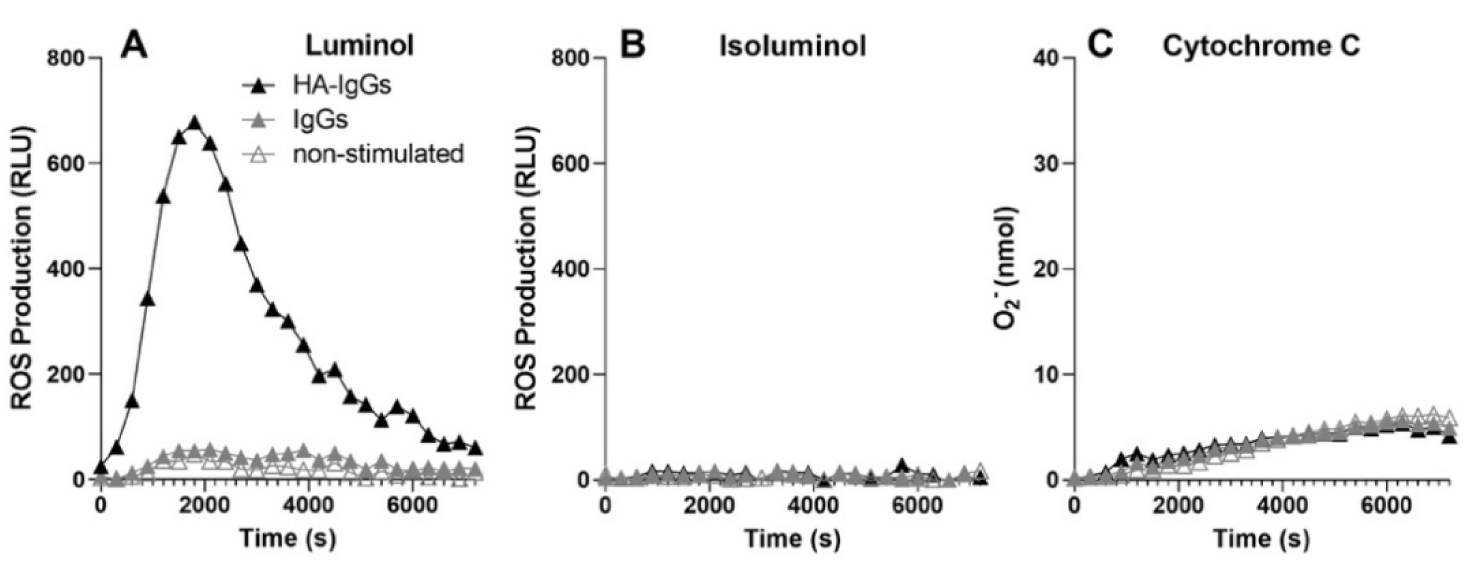
Human neutrophils stimulated with HA-IgGs produce intracellular reactive oxygen species. Neutrophils were incubated in indicated conditions, in the presence of either **A)** 10 µM luminol, **B)** 10 µM isoluminol, or **C)** 2.5% cytochrome C (v/v). ROS production was measured (37°C) simultaneously by luminescence intensity (luminol and isoluminol) and optical density (cytochrome C) every 5 min and for the indicated times. Results are from one experiment, representative of n=4 experiments performed in identical conditions with neutrophils from different donors.

### FcgRI signaling is necessary for ROS production by HA-IgG-stimulated neutrophils

The FcgRIIIb-specific shedding enzyme PI-PLC diminished ROS production elicited by HA-IgGs, by approximately 35% (**Fig. 5A**). Antibodies against FcgRIIa (IV.3), FcgRIII (3G8), FcgRIIIb (11.5), or Fc-Block, each prevented ROS production by more than 80%. Similarly, pre-incubating neutrophils with the 10.1 antibody blocked ROS production, showing the involvement of FcgRI also in this cellular function. While the up-regulation of FcgRI involved FcgRIIa, ROS production appeared to necessitate the engagement of all FcgRs monitored: FcgRI, FcgRIIa, FcgRIIIa, and FcgRIIIb. Inhibition of Syk, PI3K pathways, all but obliterated ROS production (**Fig. 5B**). Blockades of PLC, PKC, or MEK pathways also partly diminished ROS production. Nexinhib20 prevented ROS production, in a concentration-dependent fashion, but with higher potency than for FcgRI expression (**Fig. 5C**). Taken together, these results show the necessity of many pathways, including FcgRI signaling, for optimal ROS production.

**Figure 5.**
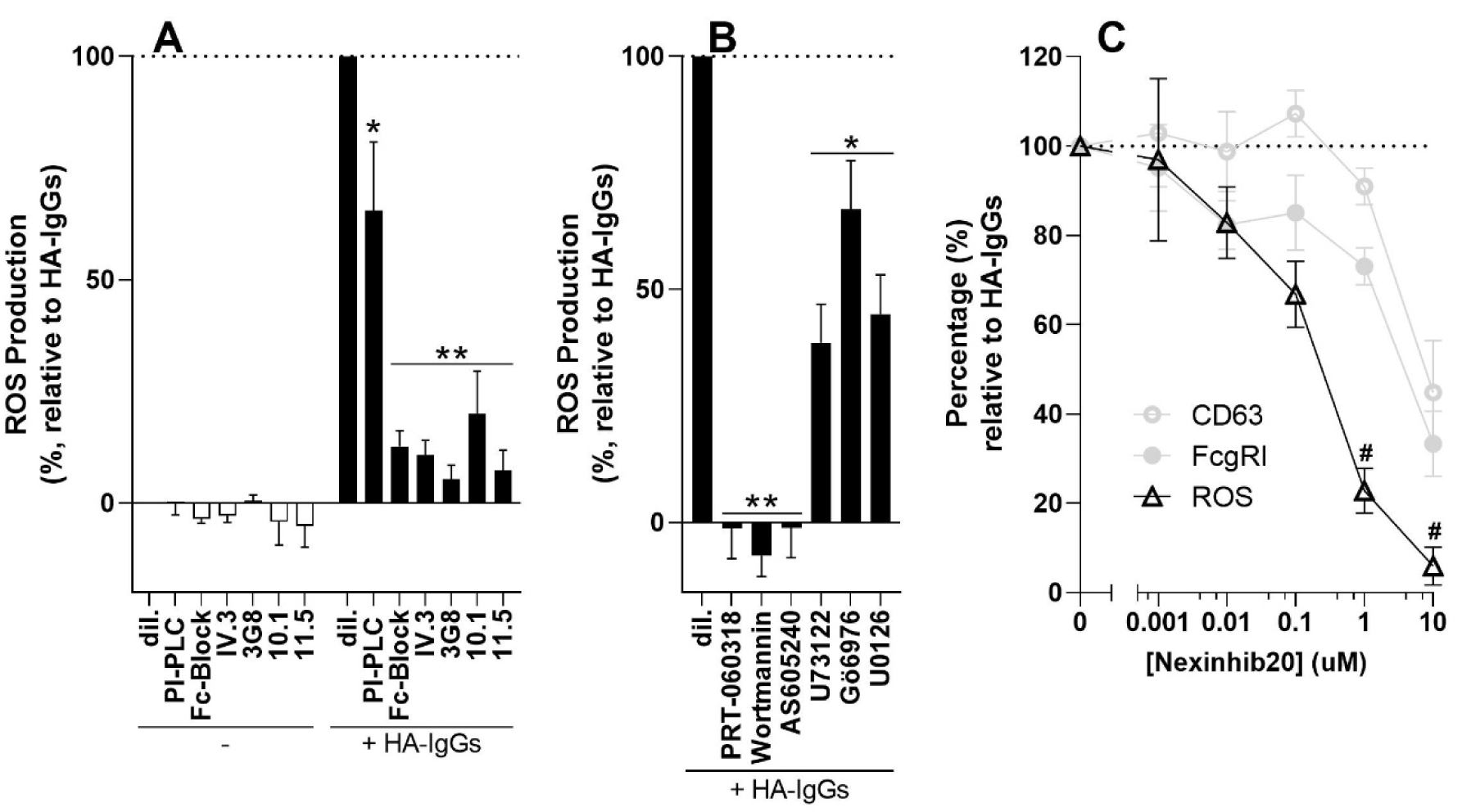
FcgRI activation contributes to the production of reactive oxygen species. **A)** Neutrophils were incubated with **(A)** PI-PLC, Fc-Block, indicated antibodies, or **(B)** with indicated pharmacological inhibitors, or **C)** with indicated concentrations of Nexinhib20 for 15 min, before stimulation with HA-IgGs. ROS production was measured by luminescence intensity (luminol). Results are expressed as the percentage of ROS production, relative to HA-IgGs alone (dil., black bar). Mean ± SEM, n=4. *Significantly lower than HA-IgGs alone. *p<0.05, **p<0.0001. ^#^Significantlly lower than CD63 and FcgRI (Panel C: pale gray data was reproduced from Fig. 3C, for easier comparison).

### Bacterial components magnify FcgRI-dependent ROS production

Given the potentiation of FcgRI expression by fMLP and LPS, we activated neutrophils with these, and other agonists, and measured ROS production. While having no effect by themselves, fMLP and LPS each had an enormous amplifying impact on the ROS production elicited by HA-IgGs up to 6-fold (**Fig. 6A**). For LPS, a TLR4 ligand, as little as 1 ng/ml was sufficient to increase ROS production (**Fig. 6B**). Blocking FcgRI, or addition of Nexinhib20, obliterated the effects of LPS and fMLP, reducing the production of ROS by at least 90%. Together, these results identify conditions that can be encountered by circulating neutrophils, combining the up-regulation of FcgRI and engagement of TLR4 or fMLP receptors, leading to massive ROS production.

**Figure 6.**
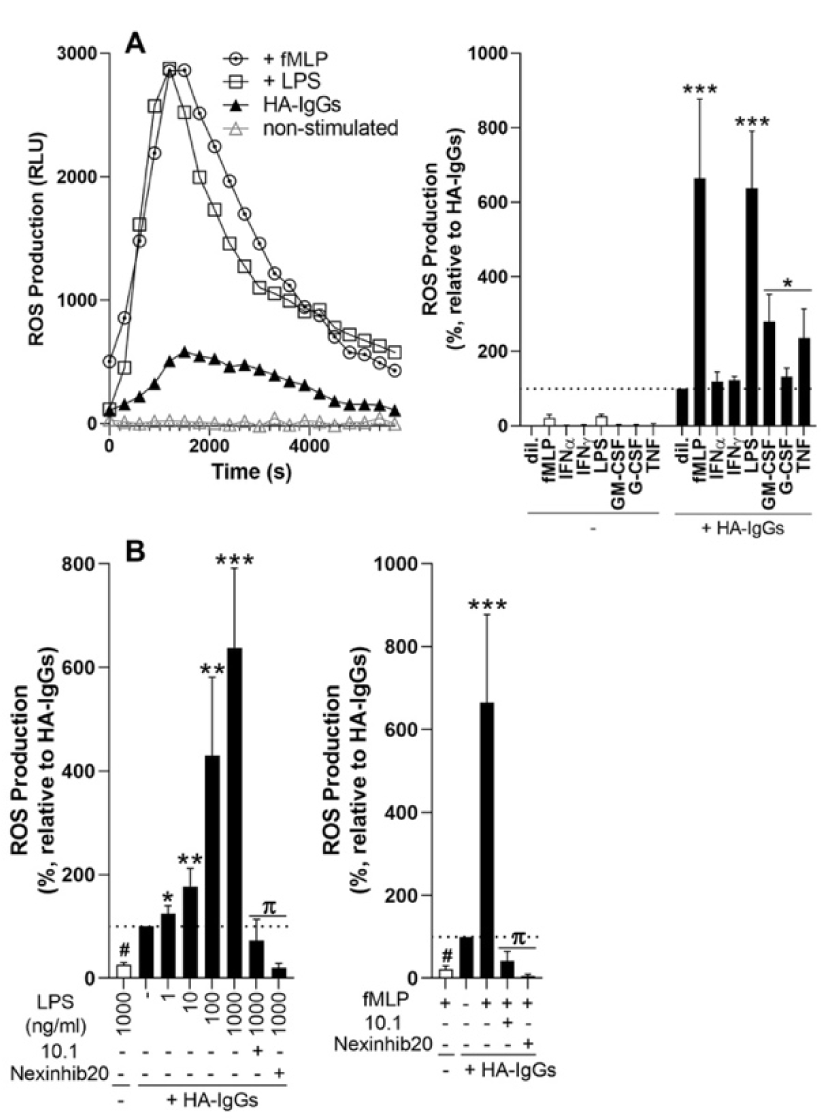
Combinations of fMLP or LPS with HA-IgGs magnify ROS production. Neutrophils were incubated with indicated agonists for 10 min (37°C) before stimulation with HA-IgGs. ROS production was determined. **A)** Typical traces of combinations with fMLP and LPS, representative of n=3 experiments, are shown. **B)** Neutrophils were incubated with indicated concentrations of LPS, or with fMLP (100 nM), then stimulated with HA-IgGs. Bar graphs: Results are expressed as the percentage of ROS production relative to HA-IgGs alone (dil., black bar) and are the mean ± SEM, n=3. Significantly lower^#^ or higher* than HA-IgGs alone (*p<0.05, **p<0.01, ***p<0.001). Significantly lower^π^ than HA-IgGs+LPS (or +fMLP) p<0.01.

Schematics, summarizing the main findings of the present study, are presented in **Figure 7**.

**Figure 7.**
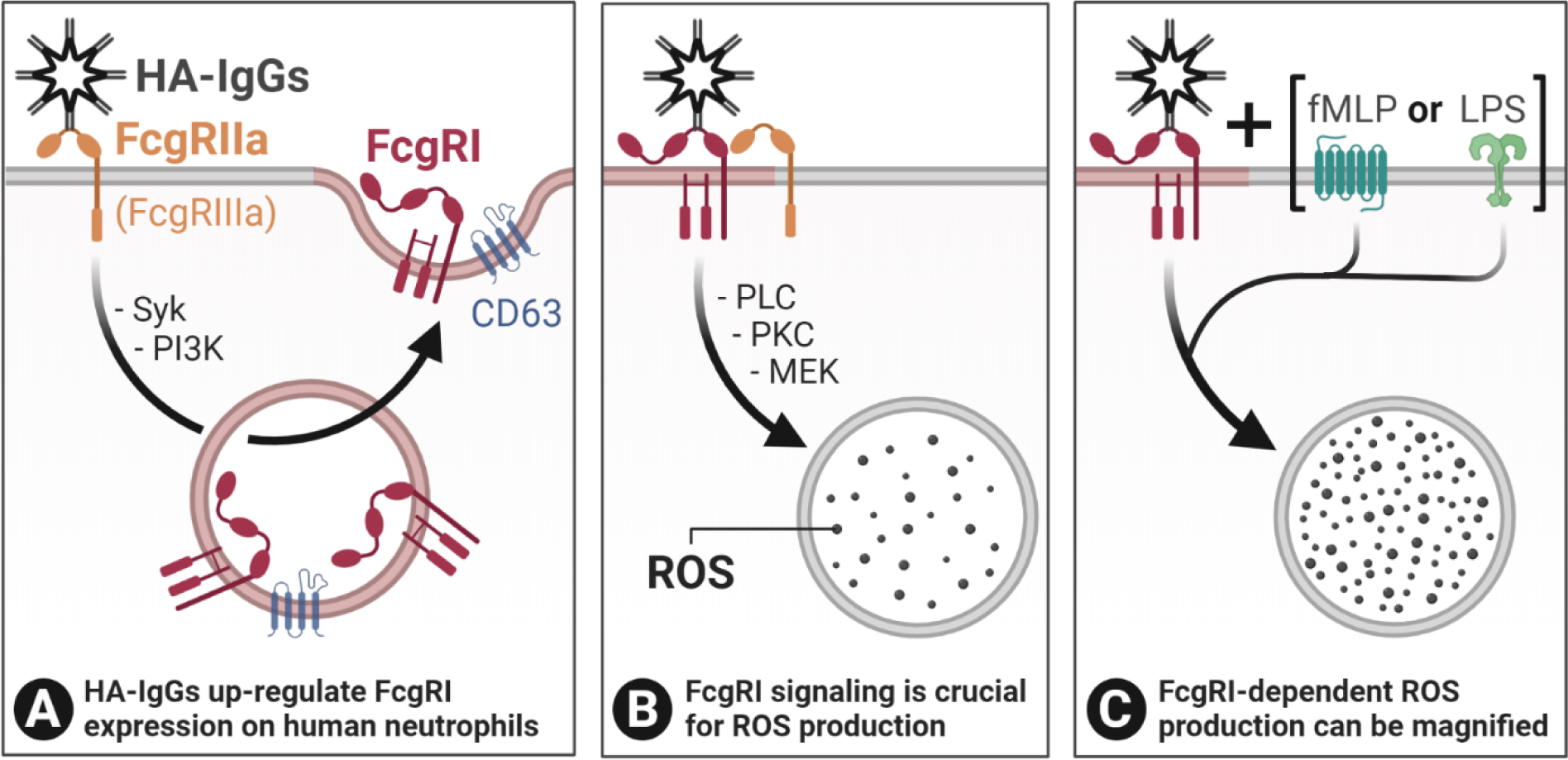
Schematics: HA-IgGs rapidly up-regulate FcgRI expression at the surface of human neutrophils in a FcgRII-dependent fashion and contribute to ROS production. **A)** Binding of HA-IgGs to FcgRIIa (and, possibly, FcgRIIIa) up-regulate FcgRI surface expression on human neutrophils, from CD63 positive, primary granules, in a Syk and PI3K-dependent fashion. **B)** Up-regulation and engagement of FcgRI by HA-IgGs is necessary for intracellular ROS production and involves PLC, PKC, and MEK. **C) In the presence of bacterial components such as LPS or fMLP**, ROS production by HA-IgGs-FcgRI complexes is potentiated. Abbreviations: HA-IgGs; heat-aggregated IgGs, fMLP; formyl-methionyl-leucyl phenylalanine, LPS; lipopolysaccharide, MEK; mitogen-activated protein (MAP)/ERK kinase, PI3K; phosphatidylinositol 3-kinase, PLC; phospholipase C, PKC; protein kinase C, ROS; reactive oxygen species, Syk; Spleen tyrosine kinase. *Created with BioRender*.*com*

## Discussion

In the present study, HA-IgGs rapidly up-regulated the surface expression of FcgRI on human neutrophils by mobilizing it from intracellular stores. In this regard, HA-IgGs proved to be more potent than any of the previously reported agonists (20, 21, 51). Among the several cell responses assessed, we showed that FcgRI engagement was specifically essential for ROS production. Also, additional stimuli synergized with HA-IgGs, magnifying ROS production. These results reveal key-elements in our understanding of interactions between immune complexes and neutrophils.

The up-regulation of FcgRI by HA-IgGs was dependent on FcgRIIa engagement and activation of Syk and PI3K, in line with an FcgR-dependent event (52, 53). None of the inhibitors blocked FgcRI up-regulation by more than half, suggesting that several pathways work parallel, leading to degranulation. PLC, PKC, GSK-3, and MEK pathways, also downstream of ITAM signaling and leading to Ca^2+^ elevation and gene expression (54), were unnecessary, indicating that FcgRI exocytosis is regulated by earlier events, upstream of those pathways.

In the present conditions, FcgRI up-regulation occurred within seconds, without the need for *de novo* protein synthesis, which points towards different mechanisms than in previous studies where patients or neutrophils were exposed to agonists for several hours, or days (20, 21, 51). Pre-formed FcgRI appears to be localized in intracellular granules, as suggested by the use of Nexinhib20, an inhibitor of granule exocytosis (55), which efficiently prevented the up-regulation of FcgRI. Moreover, hand-in-hand up-regulation of FcgRI and CD63, suggesting that primary granules may harbor FcgRI. Specialized imaging techniques will help in clarifying its precise localization. Also, the concealment of FcgRI inside resting cells may constitute a checkpoint that prevents neutrophils in circulation from getting unduly stimulated; rapid exposure of this high-affinity receptor releases fuller responsiveness when conditions require it, upon interaction with immobilized immune complexes, for example.

Complexed-IgGs have long been known to bind to neutrophils through FcgRs and stimulate cellular responses, including ROS production (56). Both FcgRIIa and FcgRIIIb have been implicated in this process, often by using blocking antibodies IV.3 and 3G8, respectively (39). Using similar and additional tools, our results around the expression of FcgRI also supported the involvement of FcgRIIa, but not that of FcgRIIIb. The FcgRIIIb-specific 11.5 blocking antibody, actually enhanced FcgRI up-regulation, as did PI-PLC, a FcgRIIIb-specific shedding enzyme. Moreover, the 3G8 blocking antibody (57), which does not discriminate between the two FcgRIII isoforms, did not affect FcgRI up-regulation, supporting the concept of a decoy role for FcgRIIIb (12, 58, 59) in this process. Results leave the possibility of a role for FcgRIIIa. This point may be relevant because FcgRIIIa, like FcgRI, is a classic activating receptor associating with an ITAM-signaling system; it might mediate many events previously attributed to FcgRIIIb (60). Alternatively, the balance in expression between the two isoforms might influence cellular responsiveness (60). Tools better suited for the specific study of FcgRIIIa will help in clarifying this issue.

ROS production could be prevented by blocking each of the receptors mentioned above. On the other hand, blocking FcgRI prevented ROS production elicited by HA-IgGs while it did not affect viability, degranulation, or cytokine release. This fact suggests a substantial and specific role of FgcRI in regulating ROS production. Also, inhibitors of PI3K, PLC, PKC, MEK1/2, each efficiently prevented this cell’s response. Moreover, Nexinhib20, an agent preventing Rab27a-JFC1 interactions, also inhibited the exocytosis of FcgRI, adding to its recognized anti-inflammatory properties (55). This profile of results clearly shows that many signaling pathways are to be solicited for ROS production, suggesting a tight regulation of this critical cell function.

One of the most salient findings presented here was the potentiation of HA-IgG-elicited ROS production by bacterial materials. Indeed, fMLP or LPS enhanced this cell’s response by manifold. On the one hand, these results show that the minute presence of microorganisms can substantially impact neutrophil activation and could profoundly affect the course of an inflammatory reaction, possibly leading to an overt, ill-regulated response. On the other hand, they indicate a crucial role for FcgRI in the process since preventing its engagement virtually obliterated ROS production, even in the presence of fMLP of LPS.

The present study finds particular relevance in the context of immune complex-mediated inflammatory disorders where current models include the capture of neutrophils by endothelial cells (61), on which immobilized immune complexes can elicit the rapid attachment of neutrophils (62) and feed inflammatory flares associated with human autoimmune diseases, such as glomerulonephritis, arthritis, rheumatic fever, and lupus. Neutrophils binding to deposited immune complexes in the kidneys of patients with lupus nephritis, one of the significant complications of lupus, may contribute to its pathogenesis (63-66) through inflammatory responses such as the biosynthesis of lipid mediators, antibody-dependent cell phagocytosis, MCP-1 release, and ROS production (64). While such functions are designed for host defense, disordered responses may instead lead to tissue damage (67). In the context of autoimmune diseases, even minor microbial contamination could steer neutrophil reactiveness in harmful directions. Further studies targeting FcgRI, using distinct approaches, will help mark the involvement of this receptor in neutrophil biology.

In conclusion, the present study reveals an essential role of neutrophil-FcgRI in cellular activation in response to IgG aggregates, particularly for the production of ROS. Moreover, cross-talk between FcgRI and other signaling pathways may have a profound impact on this cell’s response. Together, these results provide molecular insights into how neutrophil inflammatory responses can become deleterious.

## Author’s contributions

**SH:** Performed most experiments, analyzed the data, wrote parts of the manuscript.

**CL:** Performed some of the experiments, including all surface marker measurements.

**PF:** Designed parts of the study.

**EB:** Designed parts of the study.

**MP:** Designed the study, analyzed the data, wrote the manuscript.

## Acknowledgments

This work was funded by a grant from the Canadian Institutes of Health Research (CIHR) to MP (grant number: MOP220733). SH is the recipient of a studentship from the Fonds de Recherche du Québec-Santé (FRQS). EB is the recipient of a new investigator award from the CIHR and is a Canadian National Transplant Research Program (CNTRP) researcher. PRF is the recipient of a tier 1 Canada Research Chair on Systemic Autoimmune Rheumatic Diseases.

The authors confirm that there are no conflicts of interest.

## Nonstandard abbreviations list

7-AAD: 7-Aminoactinomycin D
ACTB: Actin beta
ADA: Adenosine deaminase
ERK: Extracellular signal-regulated kinase
FcgR: Fc gamma receptor
fMLP: Formyl-methionyl-leucyl phenylalanine
G-CSF: Granulocyte-colony stimulating factor
GM-CSF: Granulocyte-macrophage colony-stimulating factor
GPI: Glycosylphosphatidylinositol
GSK: Glycogen-synthase-kinase
HA-IgGs: Heat-aggregated IgGs
HBSS: Hank’s Balanced Salt Solution
HOCl: Hypochlorous acid
IFN: Interferon
IgG: Immunoglobulin G
IL: Interleukin
ITAM: Immunoreceptor Tyrosine-based Activation Motif
LPS: Lipopolysaccharide
MCP: Monocyte chemoattractant protein
MEK: Mitogen-activated protein/extracellular signal-regulated kinase kinase
MIP: Macrophage inflammatory protein
MMP-9: Matrix metallopeptidase 9, also known as gelatinase B
MPO: Myeloperoxidase
NET: Neutrophil extracellular trap
PI3K: Phosphatidylinositol 3-kinase
PI-PLC: Phosphatidylinositol-Specific Phospholipase
PKC: Protein kinase C
PLC: Phospholipase C
ROS: Reactive oxygen species
SEM: Standard error of the mean
Syk: Spleen tyrosine kinase
TLR4: Toll-like receptor 4
TNF: Tumor necrosis factor

