## Supplemental figures for "IgG-aggregates rapidly up-regulate FcgRI expression at the surface of human neutrophils in a FcgRII-dependent fashion: A crucial role for FcgRI in the generation of reactive oxygen species"

### Supplemental Materials

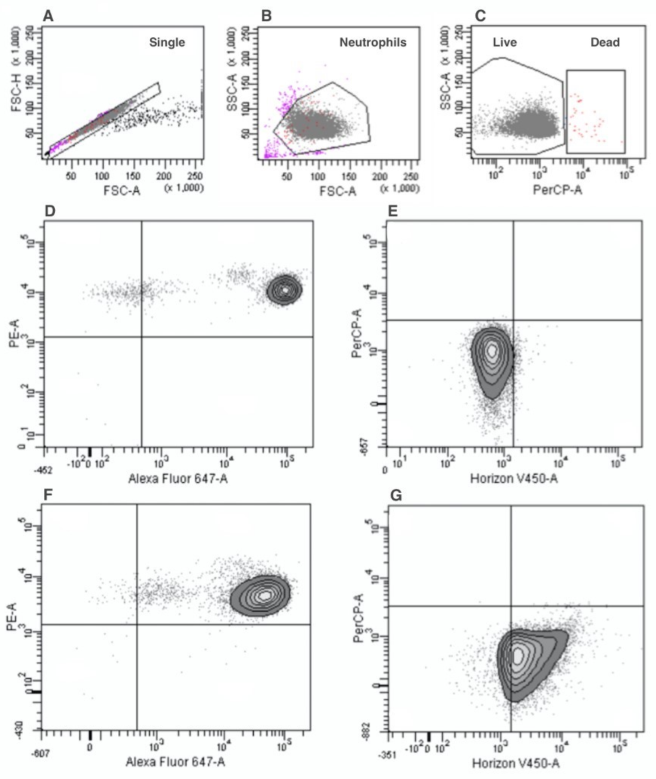


**Supplemental Figure S1**. Description of the flow-cytometric gating strategy. Single cells were targeted (**A**), neutrophils identified (**B**), and live cells were separated from dead cells (**C**). Representative graph of neutrophils stained with PE-anti-human CD32, AlexaFluor 647-anti-human CD16 and V450-anti-human CD64 in non-stimulated (**D, E**) and HA-IgGs conditions (**F, G**). Abbreviations: FCS: Forward Scatter; SSC: Side Scatter.

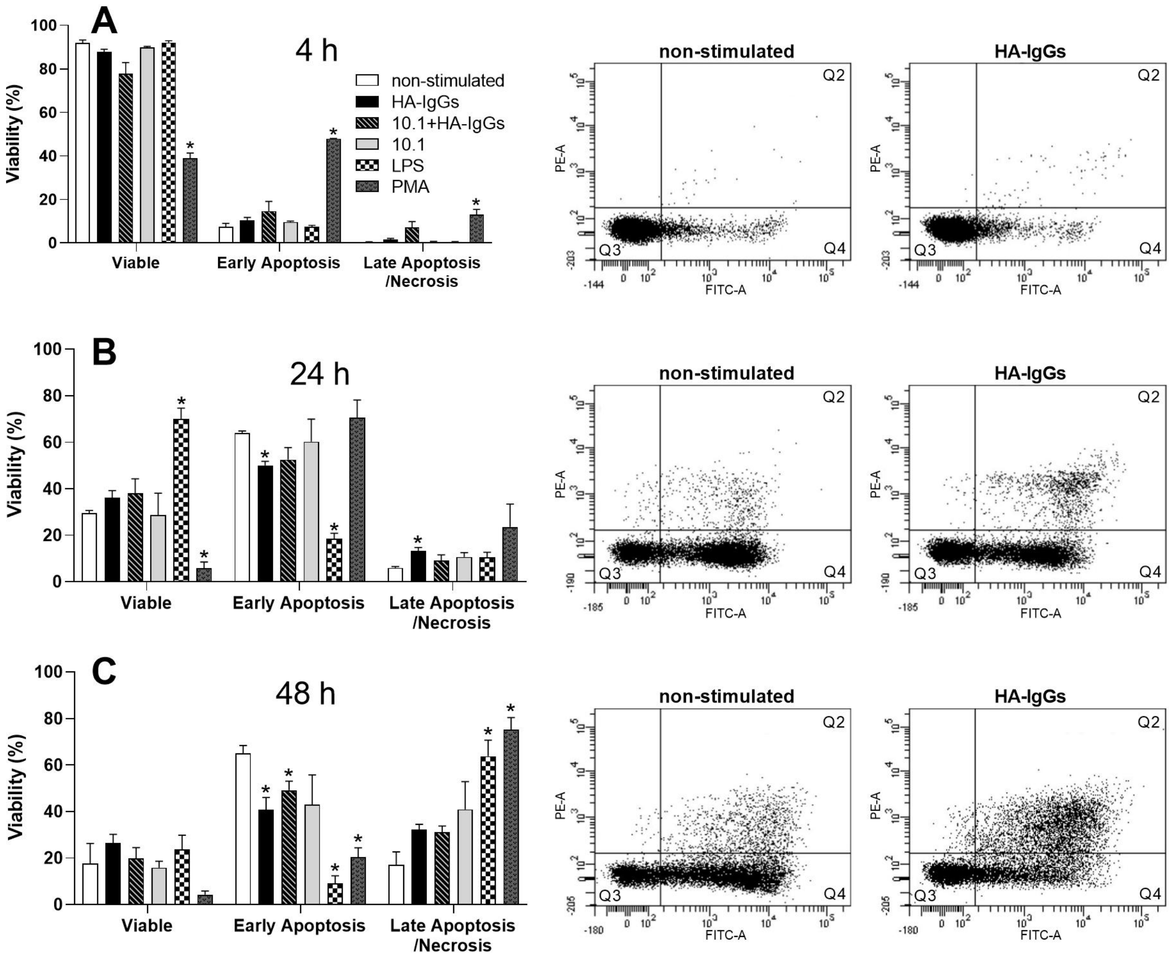


**Supplemental Figure S2. Impact of HA-IgGs on neutrophil viability.** Human neutrophils were stimulated with HA-IgGs, or indicated agents for up to 48 hrs, and cell viability was measured by flow cytometry, as described in *Methods*. *Internal controls*: LPS was used as an apoptosis-delaying, necrosis-inducing agent (68); PMA as an efficient reducer of cell viability (69). HA-IgGs: 1 mg/ml; LPS: 1 μg/ml; PMA: 10 nM. Results are the mean ± SEM, n=3. *Significantly different, compared to non-stimulated cells, $p<0.05$. Typical results from one experiment for non-stimulated and HA-IgG-stimulated cells are shown on the right (Q3 = Viable; Q4 = Early apoptosis; Q2 = Late apoptosis/necrosis).

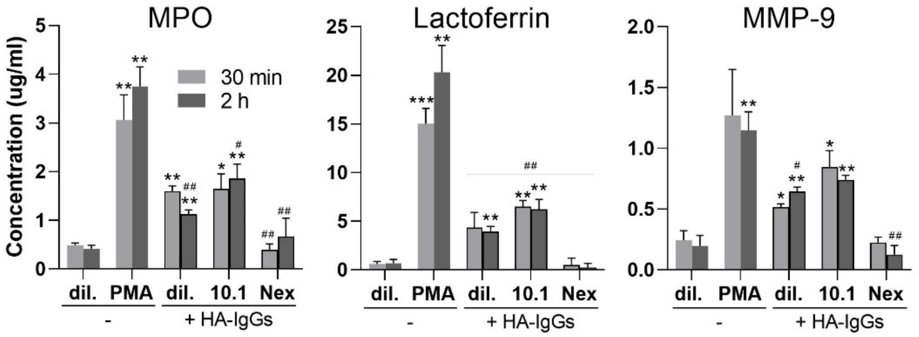


**Supplemental Figure S3**. **Human neutrophils stimulated with HA-IgGs release intracellular granules.** Following indicated stimulations, cell-free supernatants were analyzed for their content in myeloperoxidase (MPO; primary granules), lactoferrin (secondary granules), and gelatinase (MMP-9; tertiary granules) with commercially-available ELISAs, according to the manufacturers' instructions. *Positive control:* PMA was used as an efficient inducer of exocytosis (degranulation). HA-IgGs: 1 mg/ml; PMA: 10 nM; Nex: 10 µM. Results shown are the mean ± SEM, n=3. *Significantly different, compared to non-stimulated cells (dil.), or #compared to PMA. *$p<0.05$, **$p<0.01$, ***$p<0.001$.

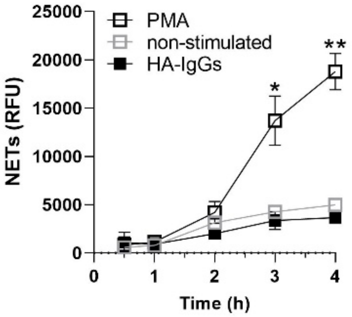


**Supplemental Figure S4. Absence of NET formation by HA-IgG-stimulated human neutrophils.** Human neutrophils were incubated at 37°C (5% $CO_2$) with HA-IgGs (1 mg/ml) or PMA (10 nM) for up to 4 hrs. NET induction was assessed by the addition of SYTOX green (5 μM) and subsequent fluorescence reading (excitation/emission: 504/523 nm), as described in *Methods*. Results are expressed as relative fluorescence units (RFU). Mean ± SEM, n=3. *p<0.05, **p<0.0001

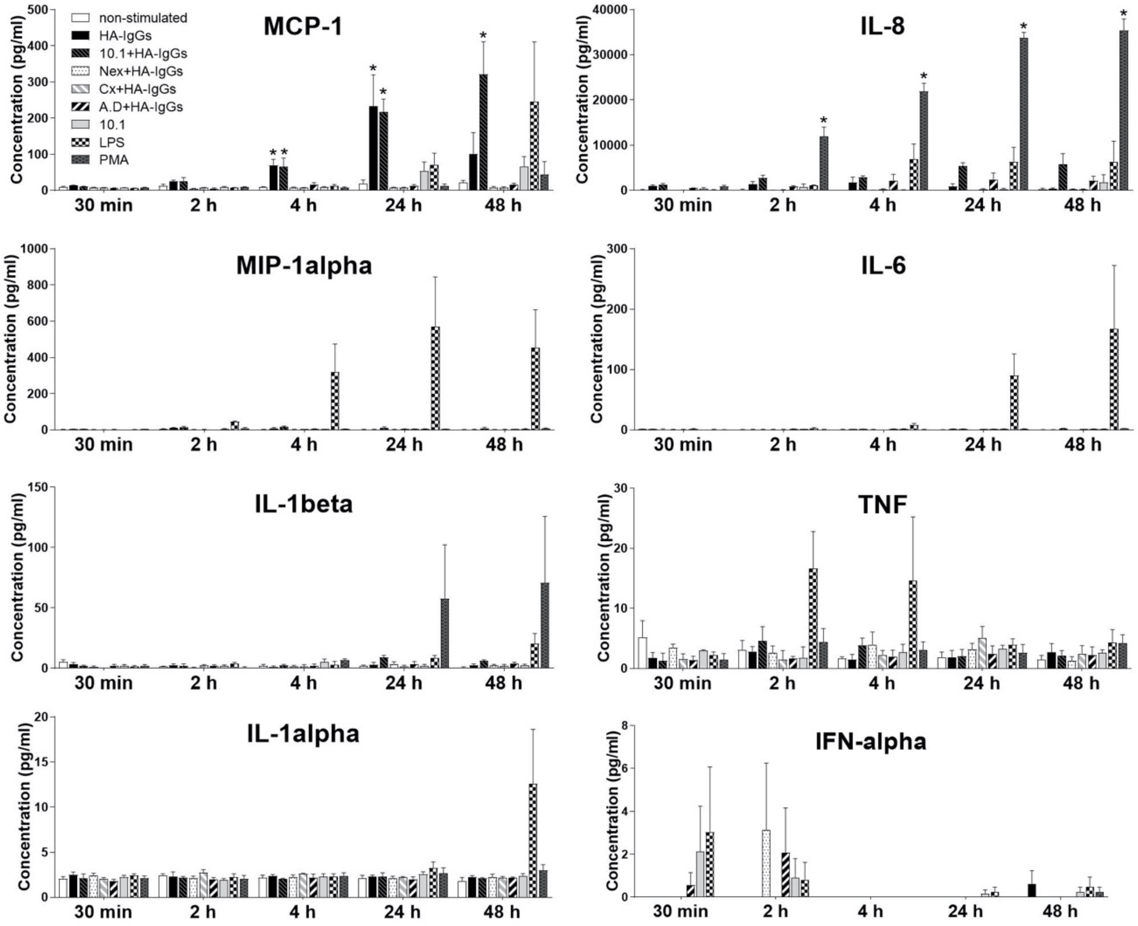


**Supplemental Figure S5**. **Stimulation of human neutrophils with HA-IgGs induces MCP-1 release.** Following indicated incubations, cell-free supernatants were analyzed for chemokine/cytokine content at indicated times. CCL2 (MCP-1), CXCL8 (IL-8), CCL3 (MIP-1alpha), IL-6, IL-1beta, TNF, IL-1alpha, and IFN-alpha were monitored. LPS and PMA were used as positive controls for the release of cytokines. HA-IgGs: 1 mg/ml; LPS: 1 µg/ml; PMA: 10 nM. Results shown are the mean ± SEM, n=3. *Significantly higher than non-stimulated, $p<0.05$. Nex: nexhinib20, Cx: cycloheximide, AD: actinomycin D.
